# Local adaptations in wing-pattern and life history trait plasticity in a butterfly: humidity as a cue where temperature is unreliable

**DOI:** 10.1101/2025.10.15.682538

**Authors:** Indukala Prasannakumar, Freerk Molleman, Urszula Walczak, Ullasa Kodandaramaiah

## Abstract

Many butterflies have wet and dry season morphs with large and small wing eyespots respectively. Eyespot size plasticity is adaptive because the morphs function to avoid predation in their respective season. Eyespot size has been shown to be regulated by rearing temperature in many species. However, temperature is unreliable in some regions because it poorly predicts seasons, and other cues such as humidity may be more reliable. We investigated inter-population differences in cue use for eyespot plasticity in the butterfly *Melanitis leda*. We reared butterflies from three Indian populations under combinations of temperature and humidity. Butterflies from a population (Vithura) where humidity differentiates seasons but temperature does not, responded only to humidity. Butterflies from a population (Tirunelveli) where seasonality is not distinct, and where temperature and humidity are both unreliable, also responded only to humidity. Butterflies from another population (Coimbatore) where seasonality is not distinct, but where temperature has the highest intra-annual variation, responded only to temperature. This suggests local adaptation in cue use. Life-history traits also differed among populations, with the two populations from more arid regions developing faster and attaining larger body sizes than the one from the humid region. Fast development may be adaptive in dry regions where suitable host plants are available only briefly, while large body size may confer desiccation resistance. We show for the first time that humidity can regulate eyespot size, and that reaction norms vary across populations, fitting to regional climates.

## INTRODUCTION

Widespread species face diverse selective pressures because they inhabit multiple geographic regions and habitats that vary in environmental conditions ^1–3^. Major environmental variables that differ across populations of a widespread species include climatic factors such as temperature ^4^, precipitation ^5^, humidity^6^, and photoperiod^7^. In addition to these climatic factors, other variables such as presence of predators or competitors^8^ and availability of resources^9^ can also vary significantly across regions. These environmental variables play a major role in shaping an organism’s survival, growth and reproduction. Colonization of new regions typically begins with dispersal, followed by survival in the novel environment^10^. However, the persistence of these newly founded populations depends on their ability to cope with the novel environmental challenges of the new habitat^11^.

A newly founded population can cope with the novel environment in two broad ways: (i) evolution of novel, genetically controlled traits, or (ii) by utilizing existing phenotypic plasticity^12,13^. Thus, widespread species can colonize new geographic regions due to their ability to adapt to local conditions, a process known as local adaptation^14,15^. However, the evolution of novel traits is a long-term process involving allelic changes^16^. In contrast, phenotypic plasticity allows the expression of multiple phenotypes depending on the environment in which individuals develop, providing a greater degree of flexibility^17,18^. Plasticity is advantageous because it allows for immediate responses to environmental challenges, without requiring generational change^23–2719,20^. This flexibility is particularly advantageous during the early stage of colonization, when rapid responses to unfamiliar environments are crucial. While phenotypic plasticity *per se* may allow persistence in a novel geographic region in the short term, it is itself a trait with genetic underpinning that can respond to selection. As populations of a widespread species adapt locally, their reaction norms may shift over time, fitting better with the demands of the new environment^21^. Plastic traits are predicted to evolve sensitivity not necessarily to the selective factors themselves, but to indirect cues that reliably forecast those factors. As the reliability of these cues for forecasting the future environment differs between regions, populations may evolve divergent cue use. Thus, in the long run, phenotypic plasticity can become locally adapted in terms of responses to environmental cues (change of reaction norms) and the relative importance of different cues (cue use).

In butterflies, adaptations to seasonal climates often involve seasonal polyphenism in wing colorations that influence survival in the face of predation. Seasonal polyphenism in butterfly wing patterns is a well-studied example of adaptive phenotypic plasticity where individuals of the same species develop different wing patterns in response to environmental cues ^24–28^. Many tropical satyrine species have large and contrasting eyespots during the wet season, but small and inconspicuous eyespots during the dry season. These eyespots are circular markings on wings that increase survival, functioning to either intimidate predators^29,30^ or deflect attacks away from vital body parts ^31–33^. During the dry season, butterflies tend to have smaller, less conspicuous eyespots, helping in camouflage against the dry background ^34–36^. Thus, eyespot plasticity serves as an excellent example of adaptive response to changing environmental conditions.

Studies over several decades, and across multiple species, have sought to understand the developmental control of eyespot size. The overwhelming majority of these studies have highlighted the role of temperature - adult eyespots are smaller when larvae develop under cooler conditions (e.g. ^24,25,28,34,37–40^). This temperature-mediated eyespot size plasticity is adaptive in habitats where the dry season is cooler than the wet season. However, seasonal temperature regimes vary widely across geographic regions^4^. Thus, the relationship between environmental variables such as temperature and rainfall also varies across regions. In some regions, temperature is not a reliable predictor of rainfall. Thus, having a conserved reaction norm across all populations could lead to ill-fitting phenotypes. Evolution is, therefore, expected to favor local adaptation of thermal reaction norms, especially in widespread butterfly species exhibiting seasonal polyphenism. Indeed, studies comparing populations of the same species from different climates have found differences in thermal reaction norms^44,45^. An alternative cue use is expected where lower temperatures do not predict drier conditions. Thus, populations of widespread species are expected to not only diverge in their thermal reaction norms but also in terms of reliance on other cues for eyespot plasticity. Environmental cues may differ strongly in terms of their reliability for plasticity. For instance, in temperate areas, photoperiod is used as a reliable cue due to its large range of variation and correlation with seasonality^46^, whereas in the tropics, day length is hardly variable and hence not a suitable cue. Similarly, temperature should be less reliable for populations where temperature has a narrow within-year range of variation. Populations of tropical satyrines that do not rely on temperature cues to determine adult eyespot size may rely on alternative cues such as those from host plants ^39,47–49^ or relative humidity (RH), which may be used independently, or in combination with temperature.

To test for differences in cue use and reaction norms between populations, we investigated inter-population differences in eyespot plasticity in *Melanitis leda* (L.) (Satyrinae: Melanitini), a widespread species distributed in a broad range of habitats across Australasia, Asia and Africa^50^. Previous studies have shown that, like many other butterfly species, *M. leda* responds to temperature, with eyespot size being larger at higher temperatures^25,39,47^. In addition, eyespot size responds to host plant species, with on average larger eyespots when larvae develop faster^39,47^. However, it is unclear whether *M. leda* uses temperature as a primary cue, especially in populations where temperature shows little seasonal variation. To address this, we collected butterflies from three Indian populations (Fig. 1) that differ in daily average rainfall and its relationship to temperature (Fig. 2). We conducted a common garden experiment by rearing larvae from each population under three combinations of temperature and RH. This design enabled us to test the independent effect of humidity and temperature on eyespot size, and to compare these across the three populations.

**Fig. 1.**
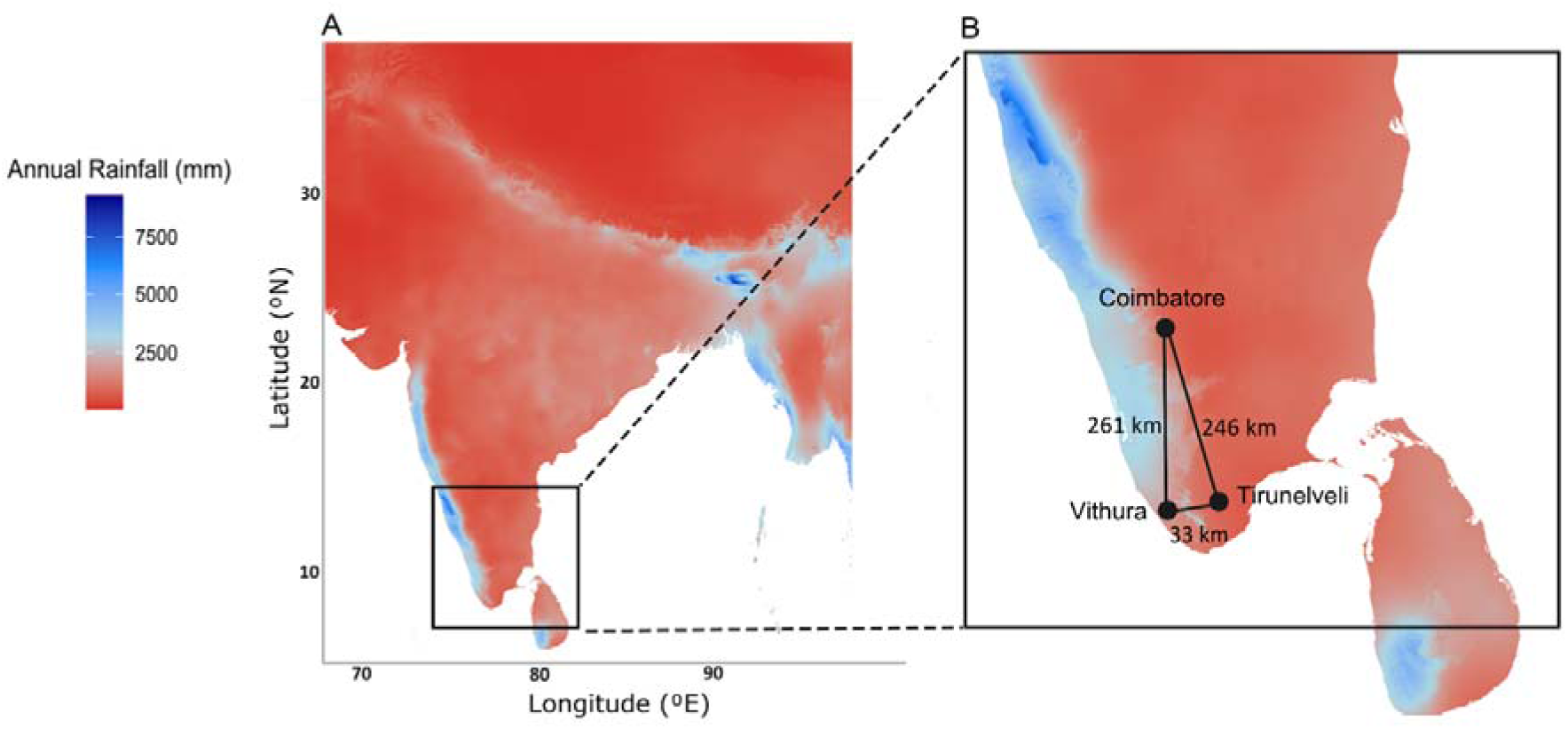
Annual rainfall distribution in the study sites in southern India. (A) Map of India showing mean annual rainfall (mm). The black square marks the region shown in panel B. (B) Enlarged view with study sites indicated by black circles: Coimbatore, Vithura and Tirunelveli. Distances between sites are shown with black lines. Rainfall is represented by a colour gradient, with darker blue indicating higher annual rainfall (Source: WorldClim^57^).

**Fig. 2.**
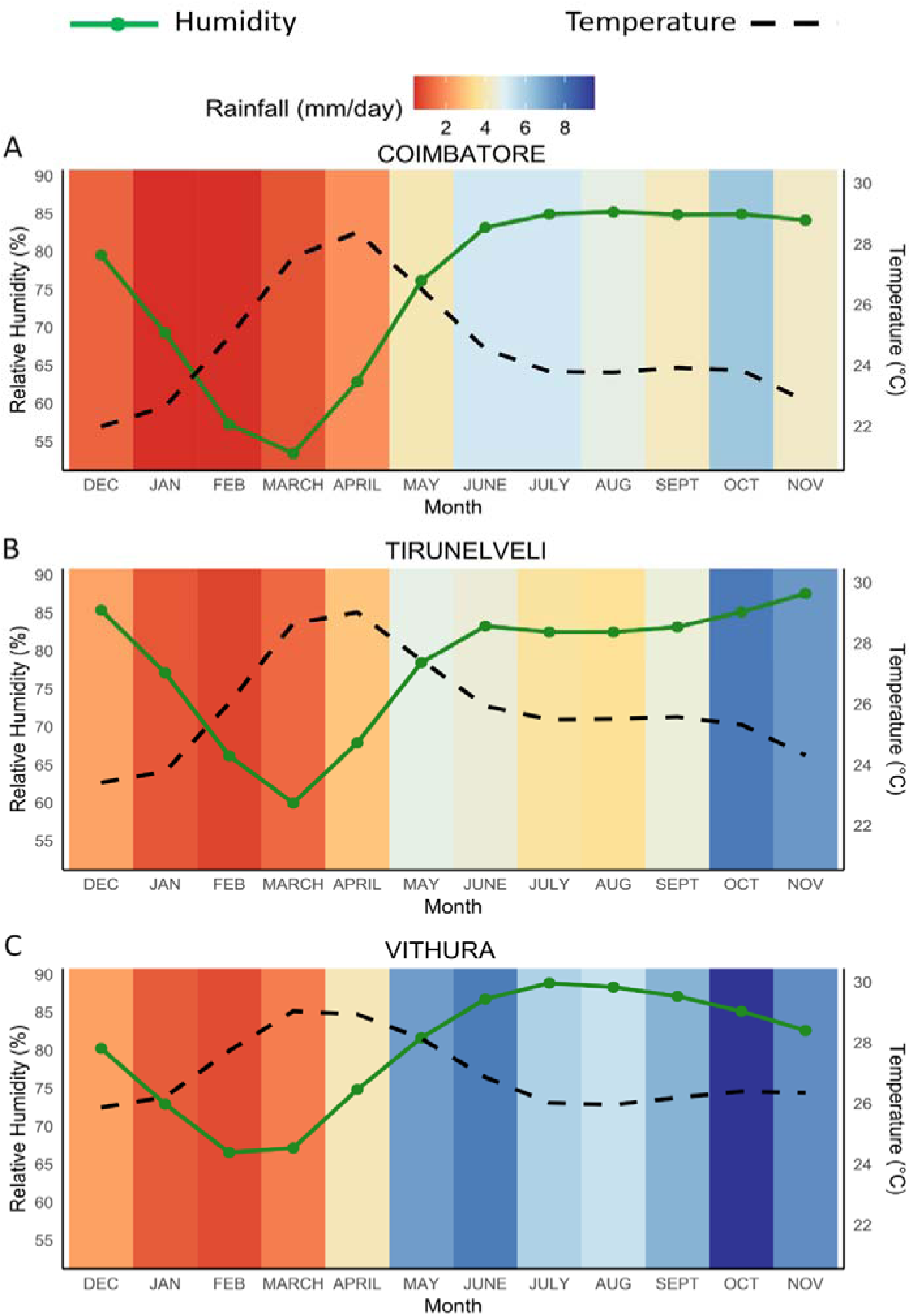
Relationship between mean monthly temperature, precipitation (mm/day) and relative humidity (%) across three locations (A) Coimbatore, (B) Tirunelveli, and (C) Vithura. Data are from 2000 to 2024 (Source: NASA POWER Data^58^).

In addition to differences in eyespot reaction norms, the populations from arid and humid regions are expected to diverge in life history traits related to desiccation tolerance. For instance, multiple studies have shown that populations and species inhabiting arid habitats have evolved larger body sizes than their counterparts in humid habitats^51–53^. Larger body size in arid conditions enhances desiccation tolerance by reducing the surface area – volume ratio ^54,55^. Moreover, larger body size can accommodate greater water and fat reserves^56^, which can help butterflies during periods of extreme drought. Furthermore, in many arid regions, the wet season tends to be short, resulting in only a short window during which high-quality host plants are available for feeding. As a consequence, populations in these environments are expected to be under strong selection for shorter developmental time to ensure completion of the life cycle before host plant resources decline in quality and quantity^42^.

We hypothesize that populations from Vithura, Tirunelveli and Coimbatore differ in their reliance on environmental cues for eyespot plasticity. While all three regions experience seasonal transitions in temperature, rainfall and humidity, the strength, timing and correlation between these cues differ (Fig. 2). In Vithura, the dry (December-April; 1.96mm average monthly rainfall) and wet (May-November; 7.12mm average monthly rainfall) seasons are strongly demarcated (Fig. 2C). Temperature does not differentiate seasons in this population. For e.g., both dry and wet seasons experience low (<27°C) and high (>27°C) temperatures. Because the larval duration in this species is usually less than six weeks^39,42,47,59^), the temperature experienced by larvae does not reliably predict the season in which adults emerge. In contrast, low (<80%) and high (>80%) RH levels clearly differentiate dry and wet seasons (Fig. 2C). Therefore, we predict butterflies from the Vithura population to rely on RH as the primary cue for eyespot plasticity. In Coimbatore, although the seasons are not as distinct as in Vithura, RH differentiates the dry (<80%) and wet (>80%) seasons (Fig. 2A), while temperature does not. Thus, we predict that butterflies from Coimbatore also rely on RH. In Tirunelveli, neither temperature nor RH is a good predictor of season or rainfall (Fig. 2B). Therefore, we predict butterflies from this population to use a combination of RH and temperature. We also predict butterflies from the dry zone, ie., Tirunelveli and Coimbatore (Fig. 1), to have greater pupal mass and shorter development time than those from the wet zone, i.e., Vithura (Fig. 1), as adaptations to a more arid climate with shorter rainy seasons.

## METHODS

### Butterfly collection

Around 40 adult butterflies were collected from each of the three locations: Coimbatore (11.0228° N, 77.0848° E), Tirunelveli (8.83° N, 77.372° E), and Vithura (8.6784° N, 77.115°E) (Fig. 1), using traps baited with fermented banana. The distance between these locations varied between 33 and 268km.Butterflies were released into population-specific cages (45 cm × 40 cm × 40 cm) in Vithura, containing two-week-old maize (*Zea mays* L.) plants for oviposition. Ripe bananas were provided as adult food. Cages were monitored daily for eggs, and newly hatched larvae were used for the experiments.

### Experimental setup

We performed a common garden experiment in which we exposed immatures of each of the three populations to three different environments. This was designed to obtain two comparisons for each population: two temperatures at the same RH, and two different RH levels at one temperature. Three insect growth chambers, all Percival Model E-36VL (Percival Scientific, Perry, USA), were set at a 12:12 Light:Day photoperiod and at the different temperature and humidity conditions: i) 27°C and 85% RH (hereafter, *27°C – 85%RH*), ii) 27°C and 60% RH (hereafter, *27°C – 60%RH*), iii) 25°C and 85% RH (hereafter, *27°C – 85%RH*). These temperatures were chosen as previous experiments showed that at these temperatures they tend to produce a mix of wet and dry-season phenotypes, and it is thus at these intermediate temperatures that we may have the best chance to detect differences between populations in their response to temperature, compared to more extreme temperatures that tend to produce only wet or only dry-season forms^47^. The three treatments were periodically rotated (i.e., reassigned) among the growth chambers to minimize growth-chamber effects.

Newly-hatched larvae were transferred onto two-week-old potted maize plants placed within nylon mesh sleeves (0.135m x 0.28m x 0.95m) and assigned to one of the treatments, and the date of larval transfer was noted. The plants were replaced every alternate day to ensure a consistent supply of fresh plants. Each sleeve contained fifteen larvae that hatched on the same day. Sleeve positions within the growth chambers were randomized regularly. Larvae were checked daily for pupation during the late larval stages. Newly-formed pupae were weighed within 48 hours after pupation to the nearest 0.0001 g using a semi-microbalance (Mettler Toledo JB1603-C), sexed by examining the genital slit of pupae ^39^. They were then transferred into 100 ml cylindrical plastic jars covered with mesh cloth for aeration so that pupae also experienced the set RH. Each jar contained one pupa and was given a unique code. The jars were then placed inside respective growth chambers until adult emergence. Thus, the egg-hatching date, pupation date, pupal mass, sex and eclosion date were recorded for each individual, along with source population and rearing conditions. Development time was calculated as the number of days from hatching to pupation.

### Eyespot measurements

Following eclosion, adult butterflies were euthanized by freezing, and their wings were carefully detached for photographing. Photographs of the separated wings were taken using a Nikon D3200 digital camera equipped with a 105mm fixed focal length lens under uniform lighting conditions within a lightbox with standardized camera settings. Across all photos, an aperture of f/18, an exposure time of 1/30 second, and an ISO sensitivity of 200 were used. Specimens were photographed against a standardized grey background to ensure consistent color calibration. Eyespot and wing measurements were taken using fixed landmarks located on the ventral surfaces of the forewings and hindwings, following protocols established in prior studies (e.g., ^28,45,47^). On the forewing, the eyespot labelled E3 (Fig. 3), and on the hindwing, the eyespot labelled E9 (Fig. 3) was used for measurement. Eyespot area was calculated from the diameter of the outer yellow ring, assuming a circular geometry, and wing area was measured by calculating the area of the red triangular region (Fig. 3). To account for individual variation in eyespot size associated with overall wing size, relative eyespot size was used for analysis, which was calculated by dividing the area of the eyespot by the proxy of the area of the wing. All morphometric analyses were performed using a custom macro in ImageJ Ver 1.52a^61^.

**Fig. 3.**
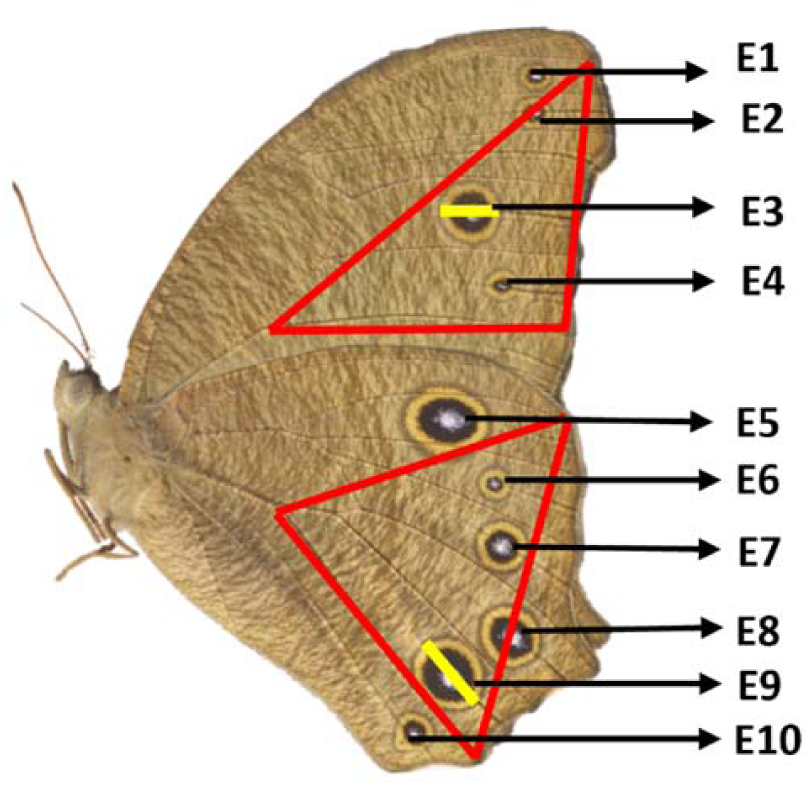
Landmarks for wing measurements for *Melanitis leda*. Area of the red triangle was used as a proxy for wing area, and the yellow line indicates the diameter of the circle used to calculate eyespot area.

### Statistical analyses

All statistical analyses were conducted using R version 4.2.1^62^ via the RStudio user interface version 2022.7.1 "Spotted Wakerobin"^63^. The effects of population and treatment on both forewing and hindwing eyespot size, as well as other life history traits such as larval development time and pupal mass, were analyzed using GAMLSS (Generalized Additive Models for Location, Scale, and Shape) through the gamlss function of the gamlss package version 5.4-3^64^. GAMLSS was used as eyespot size distribution deviated from normality, with a moderate skew and many values clustered near zero. Unlike regular Generalized Linear Models, GAMLSS allows flexible distribution fitting and can model skewness and kurtosis, providing a better fit for non-normal data. Separate analyses were conducted for males and females, because utilizing separate models reduces the number of interaction terms and facilitates a clearer interpretation of treatment and population effects within each sex^65,66^. All response variables were modelled applying a beta distribution through the BE family with the logit link function. Models were selected per the guidelines of the gamlss package^67^. The initial model included all fixed effects. Candidate models were generated by performing a model search utilizing stepGAIC’s scope option, where the null model represented the simplest model containing only the predictor variables, while the most complex model encompassed all predictor variables along with their interaction terms. The model with the lowest Generalized Akaike Information Criterion (GAIC) value was identified as the optimal model^68^. The GAIC method is a modification of the original Akaike Information Criterion^68^. The GAMLSS analyses were followed by Tukey Tests in the package emmeans Ver 1.5.2– 1^69^ for *post hoc* analysis of pairwise differences between treatments within populations.

## RESULTS

### Forewing eyespot size

There was an overall difference in eyespot size between the populations, and populations responded differently to temperature and humidity treatments: the best-fitting model with forewing eyespot size as a response variable included both main effects - treatment and population - in addition to their interaction, for both females and males (Supplementary Table S1, S2). There were also significant between-population differences in eyespot size of both sexes.

Specifically, among Coimbatore females, eyespots were larger for those reared at the lower temperature, the *25°C - 85%RH* treatment, compared to both the *27°C - 85%RH* (E=0.546, Z=3.159, P=0.0045) and the *27°C - 60%RH* (E=0.752, Z= 4.432, P<0.0001) treatments. In the other two populations, eyespot size was affected by humidity, with larger eyespots in *27°C – 85%RH* than at *27°C - 60%RH* for both Vithura (E=0.580, Z=3.668, P=0.0007) and Tirunelveli (E=0.563, Z=3.07, P=0.0061; Fig. 4) females. In males, forewing eyespot size of the Coimbatore population did not differ significantly between treatments. In the Tirunelveli population, eyespots were larger when larvae were reared at higher humidity (in *27°C - 85%RH* vs *27°C – 60%RH*: E = 0.4139, Z = 2.621, P = 0.023). Among Vithura males, eyespots were larger at higher humidity (*27°C - 85%RH* vs *27°C - 60%RH*: E = 0.784, Z = 4.859, P < 0.001), and at a higher temperature (*27°C - 85%RH* vs *25°C - 85%RH*: E = 0.5370, Z = 0.391, P = 0.002; Fig. 4).

**Fig. 4.**
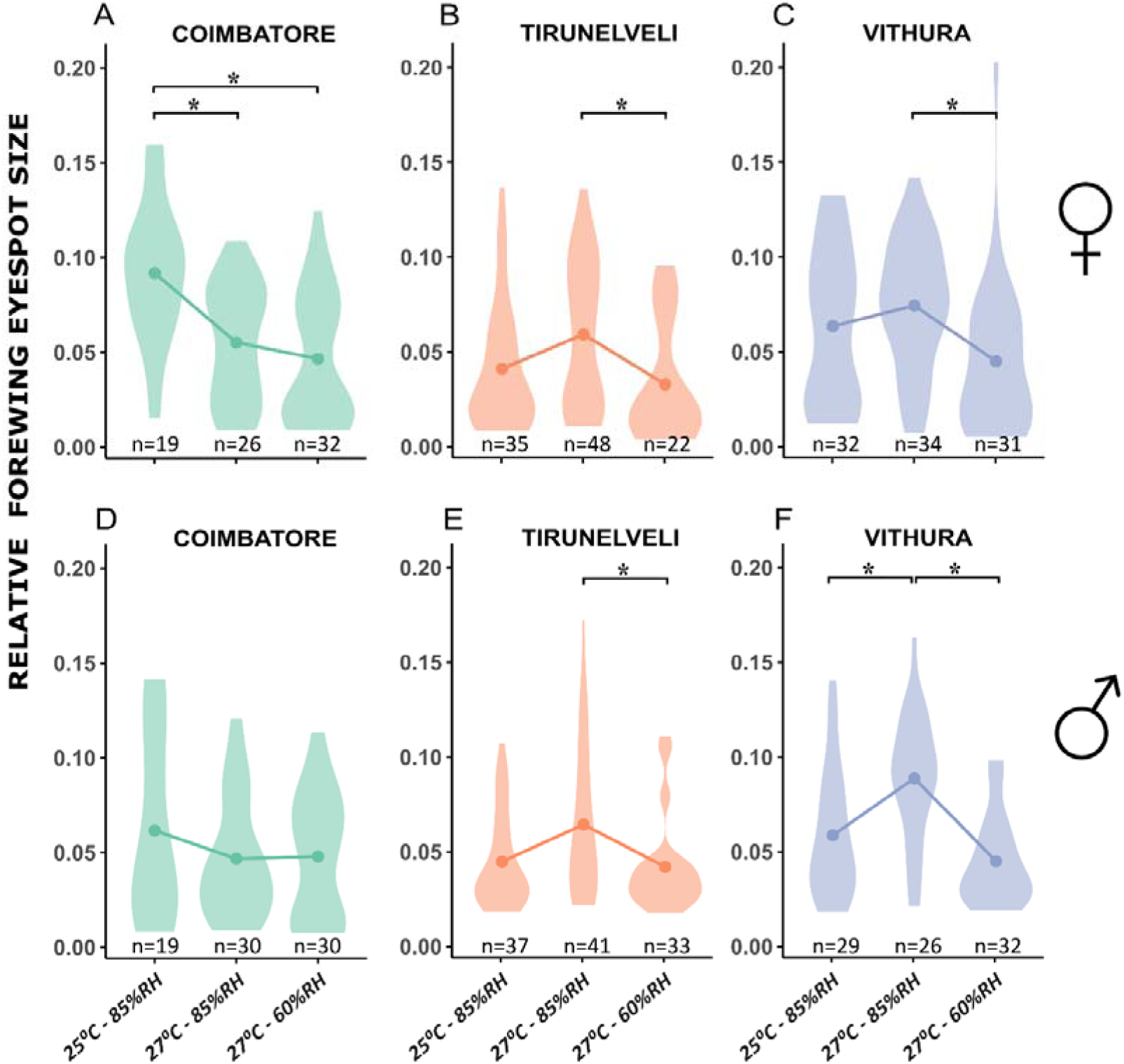
Relative forewing eyespot size across temperature and humidity treatments in *Melanitis leda* populations from Coimbatore, Tirunelveli and Vithura. Data are shown separately for females (A - C) and males (D - F). Violin plots show the distribution of eyespot sizes under three environmental conditions: *25*□*°C – 85%RH*, *27*□*°C – 85%RH*, and *27*□*°C – 60%RH*. Points indicate group means and directions of effects are indicated by connecting lines. Asterisks denote statistically significant differences between treatment groups (plJ<lJ0.05) based on *post hoc* pairwise comparisons.

### Hindwing eyespot size

Similar to that for forewing eyespots, the best-fitting model with hindwing relative eyespot size as a response variable included the main effects treatment and population, as well as their interaction effect, for both females and males (Supplementary Table S3, S4). In Coimbatore females, eyespots were larger in *25°C – 85%RH* than in both *27°C - 85%RH* (E = 0.492, Z = 2.874, *P* = 0.011) and *27°C - 60%RH* (E = 0.652, Z = 3.902, *P* < 0.001). In both Tirunelveli and Vithura females, eyespots were larger in *27°C - 85%RH* than in *27°C - 60%RH* (Vithura: E = 0.573, Z = 3.606, *P* <0.001; Tirunelveli: E = 0.578, Z = 3.123, *P*= 0.005; Fig. 5), while there was no difference in other pairwise comparisons. Relative eyespot size did not differ between treatments among Coimbatore males. Among Tirunelveli males, eyespots were larger in *27°C - 85%RH* than in *27°C - 60%RH* (E = 0.428, Z = 2.658, *P* = 0.021). Among Vithura males, eyespots were larger in *27°C - 85%RH* than in *27°C - 60%RH* (E = 0.698, Z = 4.185, *P* = 0.0001) and *25°C - 85%RH* (E = 0.39, Z = 2.405, *P* = 0.043; Fig. 5).

**Fig. 5.**
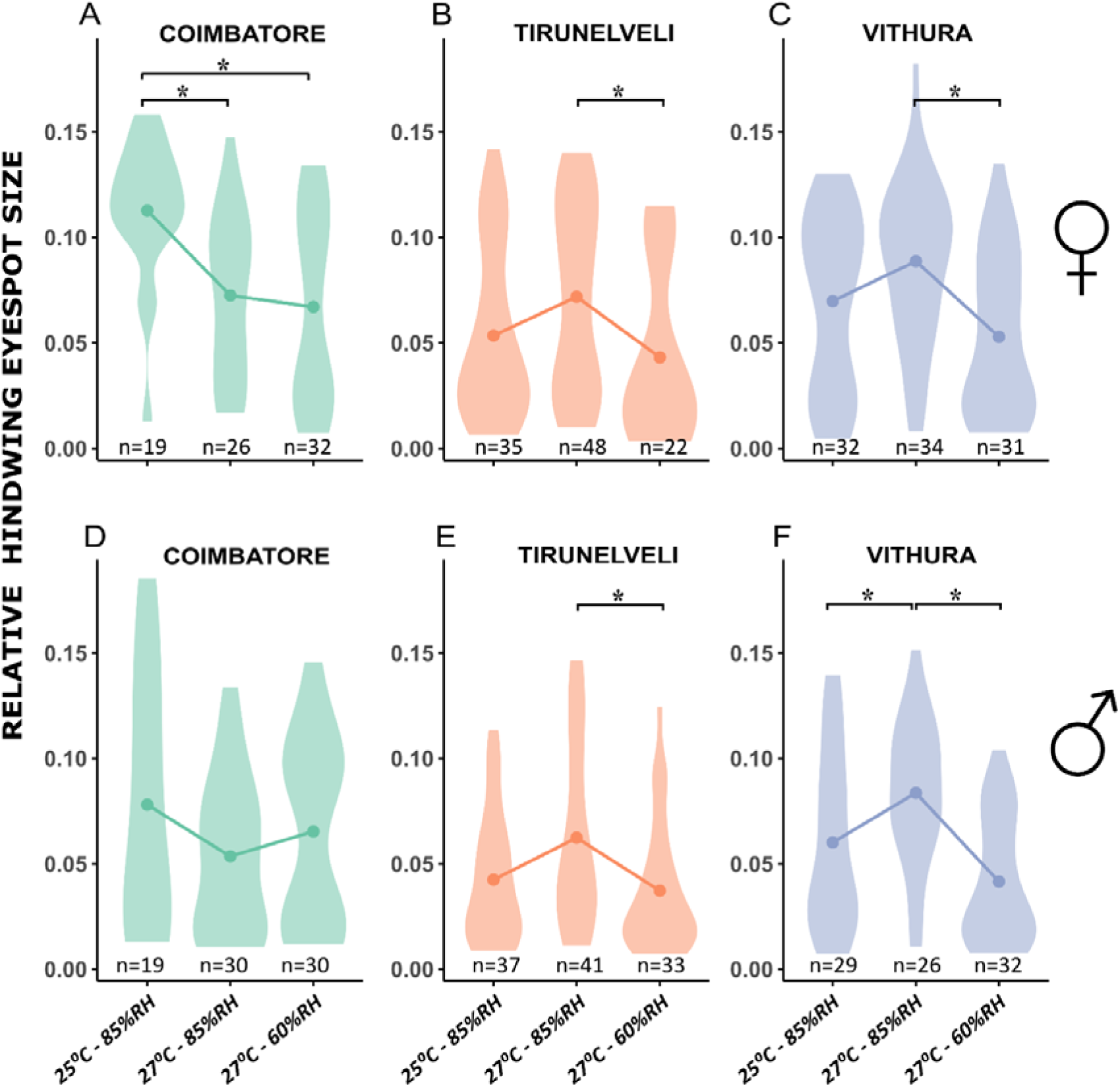
Relative hindwing eyespot size across temperature and humidity treatments in *Melanitis leda* populations from Coimbatore, Tirunelveli and Vithura. Violin plots show the distribution of eyespot sizes under three environmental conditions: *25*□*°C – 85%RH*, *27*□*°C – 85%RH*, and *27*□*°C – 60%RH*. Data are shown separately for females (A - C) and males (D - F). Points lines indicate group means. Asterisks denote statistically significant differences between treatment groups (plJ<lJ0.05 based on *post hoc* pairwise comparisons.

### Pupal mass

In females, the best-fitting model with pupal mass as a response variable retained only the independent effects of treatment and population (Supplementary Table S5). In *27°C – 60%RH*, pupae of Coimbatore females were heavier compared to those from Tirunelveli (E = 0.1777, Z = 2.380, *P* = 0.046) and Vithura (E = 0.2104, Z = 3.095, *P* = 0.006), while there was no difference between females of Tirunelveli and Vithura populations. Among males, the best-fitting model included only the effect of population (Supplementary Table S6). *Post hoc* pairwise comparisons within treatments and populations revealed that Coimbatore pupae were heavier than those from Vithura when reared at 25°C – 85%RH (E = 0.2106, Z = 2.596, *P* = 0.026). There were no other significant differences in pupal mass across treatments or populations (Fig. 6, Supplementary Fig. S1).

**Fig. 6.**
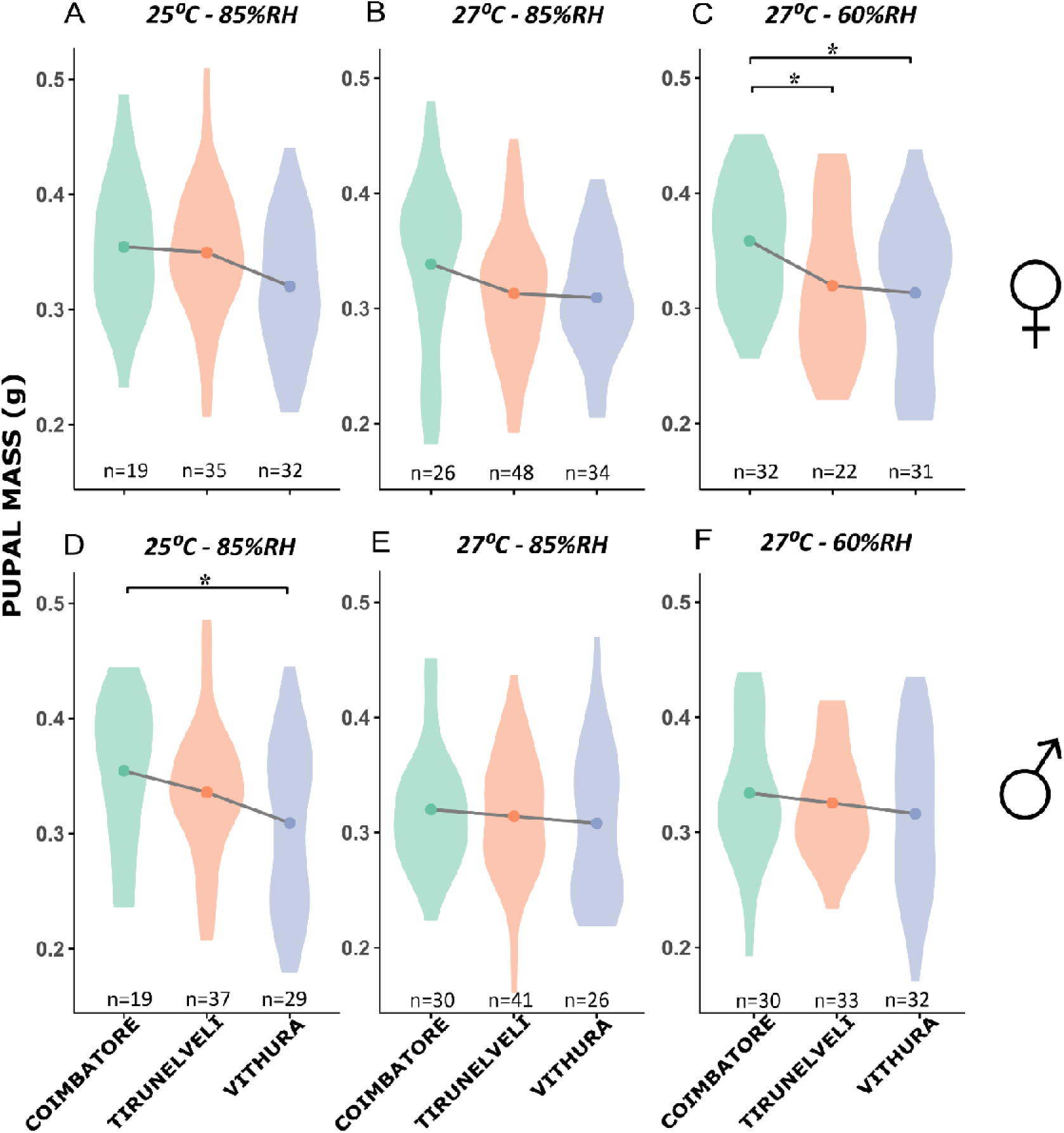
Violin plots of pupal mass (g) of butterflies from three *Melanitis leda* populations - Coimbatore, Tirunelveli and Vithura - reared under three treatments: *25*□*°C – 85%RH, 27*□*°C - 85%RH* and *27*□*°C – 60%RH*. Data are shown separately for females (A - C) and males (D - F). Each violin shows the distribution of individual larval durations (days) for a given population and treatment; the coloured point mark represents the mean. Asterisks indicate significant differences (plJ<lJ0.05) between populations based on *post hoc* pairwise comparisons.

### Development time

For both females and males, the best-fitting model included the main effects and their interaction (Supplementary Tables S7,S8). Among females, individuals from Vithura had longer larval duration than those from both Tirunelveli (E=0.27718, Z=5.508, *P*<0.0001) and Coimbatore (E=0.17456, Z=3.924, *P*=0.0003) in the *27°C - 60%RH* treatment. No other significant differences were observed between populations in the remaining treatments (Fig. 7). However, among males, there were significant differences between populations in both *27°C - 60%RH* and *25°C – 85%RH*. In *27°C – 60%RH*, larval duration was longer for Vithura males compared to Coimbatore (E=0.2564, Z=5.690, *P*<0.0001) and Tirunelveli (E=0.3288, Z=7.401, *P*<0.0001) males. Similarly, in *25°C – 85%RH*, larval duration was longer for Vithura males compared to Coimbatore (E=0.1745, Z=3.290, *P*=0.0029) and Tirunelveli (E=0.1072, Z=2.446, *P*=0.0384) (Fig. 7) males. In addition to this, some significant within-population differences across treatments were observed for both females and males (Supplementary Fig. S2).

**Fig. 7.**
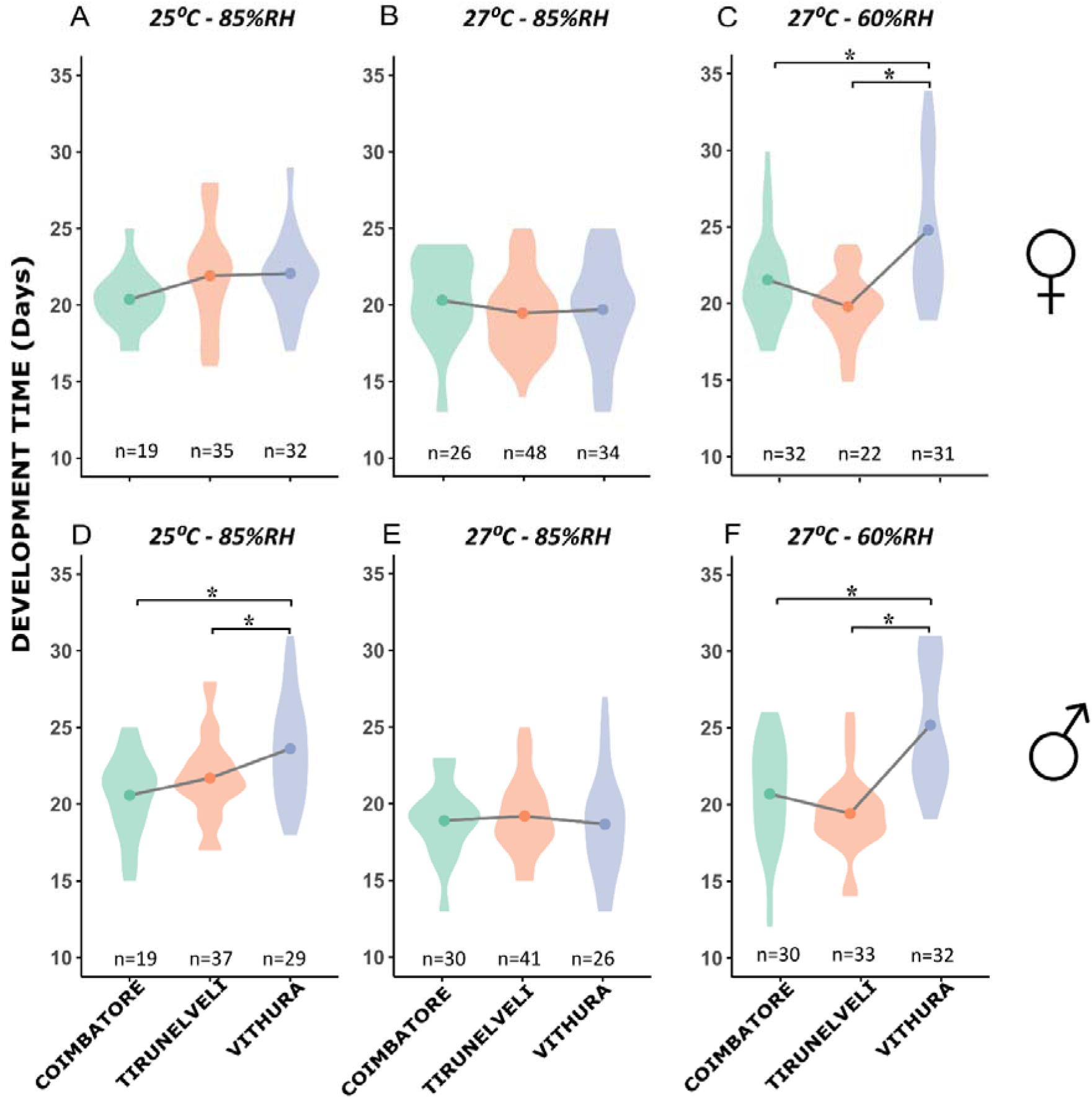
Violin plots of larval development time of butterflies from three *Melanitis leda* populations - Coimbatore, Tirunelveli and Vithura - reared under three treatments: *25*□*°C – 85%RH*, *27*□*°C – 85%RH* and *27*□*°C – 60%RH*. Data are shown separately for females (A - C) and males (D - F). Each violin shows the distribution of individual larval durations (days) for a given population and treatment; the coloured point mark represents the mean. Asterisks indicate significant differences (plJ<lJ0.05) between populations based on *post hoc* pairwise comparisons.

## DISCUSSION

We conducted a common garden experiment in which we reared three populations of *M. leda* under three conditions representing variation in temperature and humidity. We show, for the first time, that butterflies can use humidity as a cue for eyespot plasticity. In two populations, lower humidity resulted in smaller eyespots. This indicates that butterflies can sense the lower humidity during the dry season to develop into dry-season morphs. In contrast to previous studies, we found a response to temperature only in two population-sex combinations. Intriguingly, in one of these cases, the effect was in the opposite direction: higher rearing temperatures were associated with smaller eyespots, rather than larger ones. We further found that populations from drier sites had heavier pupae, even though their development time was shorter. We note that the distance between our populations (between 33 and 268km) is comparable to those used in a study^70^, that found little genetic differentiation between *M. leda* populations in southern India. Thus, there is probably significant gene flow between our study populations. Therefore, genetic differences in cue use and life history traits are less likely due to genetic drift, and more likely a result of selection that maintains these differences despite gene flow. This adds credence to our interpretation that differences between these populations are adaptations to local climate.

### Relative Humidity can mediate eyespot plasticity

Experiments on multiple tropical satyrine species across diverse regions have indicated that temperature experienced during larval development is a major driver of adult eyespot size^25,28,34,37^. This temperature-mediated plasticity is adaptive when temperature reliably predicts precipitation, i.e., when dry seasons are cooler than wet seasons. However, low temperature does not reliably predict the dry season in many regions of the tropics. Similarly, high temperature does not reliably predict the wet season in any of our three study populations (Fig. 2). Butterflies from habitats where temperature is a poor predictor of seasons are expected to rely on additional or alternative environmental cues to predict seasonal changes. While precipitation differentiates dry and wet seasons, butterflies may not be able to gauge rainfall directly. Since RH is directly related to precipitation (Fig. 2), it can be a reliable predictor of dry and wet seasons, and thus a useful cue for seasonal polyphenism. However, to our knowledge, no previous studies tested if RH can modulate eyespot size.

Our results suggest that RH can be used either as the sole cue or in combination with temperature to predict seasonal change. Future studies may unravel the mechanisms for this cue use. Butterflies may, for example, use information from a combination of cues, which is integrated in some way. While temperature and RH can directly influence butterfly development, these cues may also have an indirect influence through their effect on host plants. In particular, they may influence the nutritional quality of host plants, and this could be a cue for the larvae feeding on them^73^. For instance, host-plant nutritive quality is typically higher under humid and warm conditions^74,75^, so that host-plant quality can be a reliable predictor of season. Overall, cues such as humidity and host-plant quality need more attention in studies aiming to disentangle how seasonal polyphenism is regulated at a physiological level.

### Local adaptation in cue use

Different populations of a species experience distinct seasonal patterns of temperature, humidity and rainfall. Thus, having conserved reaction norms to RH or temperature within a species is likely to be maladaptive in some populations. Our results suggest that the two southern populations – Vithura and Tirunelveli – have evolved to rely on humidity to modulate eyespot size. In contrast, the northern population – Coimbatore – does not appear to be sensitive to RH, because neither sex responded to RH (Fig. 4 – 5). Females responded to temperature (in a direction opposite to that expected), but males did not. This suggests that this population does not rely on RH, and instead relies on temperature. We note that temperature has the highest intra-annual variation in this population.

Our results need to be treated with caution as we had only three populations and documented only partial reaction norms for two environmental factors for each of them. The range of temperatures used in this study represents only a small fraction of temperatures experienced by butterflies in their natural habitats. Previous studies with a wider range of temperatures have found overall larger eyespots in *M. leda* butterflies reared at higher temperatures in a population from Thiruvananthapuram (near Vithura), a population from Ghana^47^, and even the Vithura population, similar to most other tropical satyrines studied ^24,35,40,76^. Therefore, we do not claim that our study populations do not respond to temperature. Rather, we suggest that humidity may be at least as important in determining adult phenotype as temperature.

### Local adaptation in life history traits

Compared to the wet-zone population, the two dry-zone populations experience a more prolonged and drier dry season. We hypothesized that dry zone populations have evolved adaptations in life history traits to cope with the elevated desiccation stress. We found that pupal mass responded plastically to rearing conditions and that this response differed between populations, leading to body mass differences between populations under particular conditions. We found no differences in body size between low and high RH (*27°C - 85%RH* versus *27°C - 60%RH*; Supplementary Fig. S1). However, we found evidence of genetic differentiation in body size across populations. Females of the dry zone populations had higher pupal mass than their wet zone counterparts under conditions with the strongest desiccation stress, i.e., *27°C - 60%RH,* corroborating other studies showing insect populations in arid habitats to have larger body size^51,52^. There was, however, no inter-population difference at high humidity (*27°C - 85%RH*), suggesting that butterflies do not prioritize attaining large body size when desiccation stress is low. Interestingly, there was no inter-population differentiation in body size in males across the low RH treatments. This may be because large body size is more critical for females – females not only need to survive but also invest resources into eggs that can cope better with desiccation stress. For instance, larger females may be able to lay larger eggs that have greater viability under desiccation stress compared to small eggs^77–79^. Larger body size may also be related to starvation resistance^80^, which may be more strongly selected during dry seasons and in more arid^80^. Further studies may investigate how humidity during rearing and the climate of the natal habitat affect egg size, water- and fat content of butterflies, and contribute to these patterns in pupal mass.

Development time is another important life history adaptation for butterflies. When wet seasons are short, fresh host plants are only available for larval feeding for a short period^42,81^. Therefore, our dry-zone populations are expected to be selected for shorter development time than that of the Vithura population which has an extended wet season. Indeed, in the *27°C - 60%RH* treatment, larvae from the wet site took longer to develop than those from the two more arid sites (both females and males). These inter-population differences were also found in the *25°C - 85%RH* treatment in males, but not females. Thus, the dry zone populations appear to be able to maintain short development time across a broader range of conditions. Shorter development is expected to trade off with body size, with individuals developing faster attaining smaller size at maturity^82,83^. Surprisingly, under low humidity conditions, the population from the wetter site had a longer development time than the other populations (Fig. 7), but body size remained smaller than that of the other populations (significant for females; Fig 6), suggesting lower growth rate in the population from the wet site.

### Conclusions

While temperature-mediated eyespot plasticity in tropical satyrine butterflies has been extensively studied for several decades and across multiple species, it is unlikely that all butterflies rely solely on temperature to regulate eyespot size. We show, for the first time, that humidity can modulate eyespot size, either on its own, or in combination with temperature. Furthermore, we show that the reaction norms of eyespots to temperature and RH have diverged across populations, probably because they have been evolving under distinct climatic regimes. Under our experimental conditions, butterflies from the more northern population were sensitive only to temperature, while those from the southern populations were more sensitive to humidity than to temperature, suggesting local adaptation across populations in the eyespot reaction norms. Life history also responded plastically to humidity, with larger body sizes when reared at lower humidity, probably to increase desiccation resistance. Local adaptation in life history was evident as butterflies from the arid populations completed larval development more rapidly and attained larger body size than those from the humid population, probably reflecting shorter growth seasons and more desiccation stress. Our results show that studies of eyespot plasticity need to take humidity into account as a potential cue, and that adaptation to climate in terms of plasticity and life history traits can evolve in populations that are only weakly spatially isolated.

## Acknowledgements

We thank all Vanasiri lab members for their support and inputs throughout the project.

## Author contributions

IK, FM, UW and UK conceived and designed the study; IK and UK conducted the research. IK, FM, UW and UK analysed the data and drafted the manuscript. All authors reviewed and approved the final manuscript.

## Ethics declarations

Not applicable

## REFERENCES

1. Chevin, L.-M. & Lande, R. Adaptation to marginal habitats by evolution of increased phenotypic plasticity. J Evol Biol 24, 1462–1476 (2011).

2. Kawecki, T. J. & Ebert, D. Conceptual issues in local adaptation. Ecology Letters 7, 1225–1241 (2004).

3. Vergeer, P. & Kunin, W. E. Adaptation at range margins: common garden trials and the performance of Arabidopsis lyrata across its northwestern European range. New Phytologist 197, 989–1001 (2013).

4. Sorte, C. J. B., Jones, S. J. & Miller, L. P. Geographic variation in temperature tolerance as an indicator of potential population responses to climate change. Journal of Experimental Marine Biology and Ecology 400, 209–217 (2011).

5. O. Gutiérrez-Hernández. Geographic variation in temperature tolerance as an indicator of potential population responses to climate change. Journal of Experimental Marine Biology and Ecology 400, 209–217 (2011).

6. Bates, O. K. & Bertelsmeier, C. Climatic niche shifts in introduced species. Current Biology 31, R1252–R1266 (2021).

7. Lin, X., Fang, C., Liu, B. & Kong, F. Natural variation and artificial selection of photoperiodic flowering genes and their applications in crop adaptation. aBIOTECH 2, 156–169 (2021).

8. Corbett, J. J. & Trussell, G. C. Local and regional geographic variation in inducible defenses. Ecology 105, e4207 (2024).

9. Esmaeili, S., Schoenecker, K. A. & King, S. R. B. Resource availability and heterogeneity affect space use and resource selection of a feral ungulate. Ecosphere 15, e4939 (2024).

10. Travis, J. M. J. et al. Dispersal and species’ responses to climate change. Oikos 122, 1532–1540 (2013).

11. Laska, A. et al. Mechanisms of dispersal and colonisation in a wind-borne cereal pest, the haplodiploid wheat curl mite. Scientific Reports 12, 551 (2022).

12. Fox, R. J., Donelson, J. M., Schunter, C., Ravasi, T. & Gaitán-Espitia, J. D. Beyond buying time: the role of plasticity in phenotypic adaptation to rapid environmental change. Philosophical Transactions of the Royal Society B: Biological Sciences 374, 20180174 (2019).

13. Wang, S. P. & Althoff, D. M. Phenotypic plasticity facilitates initial colonization of a novel environment. Evolution 73, 303–316 (2019).

14. Colautti, R. I. & Lau, J. A. Contemporary evolution during invasion: evidence for differentiation, natural selection, and local adaptation. Molecular Ecology 24, 1999–2017 (2015).

15. Meek, M. H. et al. Understanding Local Adaptation to Prepare Populations for Climate Change. BioScience 73, 36–47 (2023).

16. Kinnison, M. & Hendry, A. The pace of modern life II: From rates of contemporary microevolution to pattern and process. in Genetica vol. 112/113 145–164 (2011).

17. Agrawal, A. A. Phenotypic Plasticity in the Interactions and Evolution of Species. Science 294, 321–326 (2001).

18. de la Mata, R. et al. Drivers of population differentiation in phenotypic plasticity in a temperate conifer: A 27□year study. Evol Appl 15, 1945–1962 (2022).

19. Chevin, L.-M. & Lande, R. When do adaptiave plasticity and genetic evolution prevent extinction of a densitiy-regulated population? Evolution 64, 1143–1150 (2010).

20. Snell-Rood, E. C., Megan E. Kobiela, Kristin L. Sikkink & Shephard, A. M. Mechanisms of Plastic Rescue in Novel Environments. Annual Review of Ecology, Evolution, and Systematics 49, 331–354 (2018).

21. Oomen, R. A. & Hutchings, J. A. Genomic reaction norms inform predictions of plastic and adaptive responses to climate change. Journal of Animal Ecology 91, 1073–1087 (2022).

22. Wolda, H. Seasonal cues in tropical organisms. Rainfall? Not necessarily! Oecologia 80, 437–442 (1989).

23. Chevin, L.-M. & Lande, R. Evolution of environmental cues for phenotypic plasticity. Evolution 69, 2767–2775 (2015).

24. Bhardwaj, S. et al. Origin of the mechanism of phenotypic plasticity in satyrid butterfly eyespots. eLife 9, e49544 (2020).

25. Brakefield, P. M. Tropical dry and wet season polyphenism in the butterfly Melanitis leda (Satyrinae): Phenotypic plasticity and climatic correlates. Biological Journal of the Linnean Society 31, 175–191 (1987).

26. Dongmo, M. A. K., Bonebrake, T. C., Hanna, R. & Fomena, A. Seasonal Polyphenism in Bicyclus dorothea (Lepidoptera: Nymphalidae) Across Different Habitats in Cameroon. Environmental Entomology 47, 1601–1608 (2018).

27. Morehouse, N. et al. Seasonal selection and resource dynamics in a seasonally polyphenic butterfly. Journal of evolutionary biology 26, (2012).

28. van Bergen, E. et al. Conserved patterns of integrated developmental plasticity in a group of polyphenic tropical butterflies. BMC Evolutionary Biology 17, 59 (2017).

29. Kodandaramaiah, U., Vallin, A. & Wiklund, C. Fixed eyespot display in a butterfly thwarts attacking birds. Animal Behaviour 77, 1415–1419 (2009).

30. Vallin, A., Jakobsson, S., Lind, J. & Wiklund, C. Prey survival by predator intimidation: an experimental study of peacock butterfly defence against blue tits. Proc Biol Sci 272, 1203–1207 (2005).

31. Halali, D., Krishna, A., Kodandaramaiah, U. & Molleman, F. Lizards as Predators of Butterflies: Shape of Wing Damage and Effects of Eyespots. lepi 73, 78–86 (2019).

32. Olofsson, M., Vallin, A., Jakobsson, S. & Wiklund, C. Marginal Eyespots on Butterfly Wings Deflect Bird Attacks Under Low Light Intensities with UV Wavelengths. PLOS ONE 5, e10798 (2010).

33. Prudic, K. L., Stoehr, A. M., Wasik, B. R. & Monteiro, A. Eyespots deflect predator attack increasing fitness and promoting the evolution of phenotypic plasticity. Proceedings of the Royal Society B: Biological Sciences 282, 20141531 (2015).

34. Brakefield, P. M., Kesbeke, F. & Koch, P. B. The regulation of phenotypic plasticity of eyespots in the butterfly Bicyclus anynana. Am Nat 152, 853–860 (1998).

35. Brakefield, P. M. & Reitsma, N. Phenotypic plasticity, seasonal climate and the population biology of Bicyclus butterflies (Satyridae) in Malawi. Ecological Entomology 16, 291–303 (1991).

36. Ho, S., Schachat, S., Piel, W. & Monteiro, A. Attack risk for butterflies changes with eyespot number and size. Royal Society Open Science 3, 150614 (2016).

37. Halali, S., Brakefield, P. M. & Brattström, O. Phenotypic plasticity in tropical butterflies is linked to climatic seasonality on a macroevolutionary scale. Evolution 78, 1302–1316 (2024).

38. Mateus, A. R. A. & Beldade, P. Developmental Plasticity in Butterfly Eyespot Mutants: Variation in Thermal Reaction Norms across Genotypes and Pigmentation Traits. Insects 13, 1000 (2022).

39. Prasannakumar, I., Molleman, F., Chandavarkar, D. & Kodandaramaiah, U. Development time integrates temperature and host plant cues for eyespot size in three tropical satyrine butterflies. Journal of Insect Physiology 163, 104814 (2025).

40. Windig, J. J. Seasonal polyphenism in Bicyclus safitza: a continuous reaction norm. (1991).

41. Adler, R. F. et al. Relationships between global precipitation and surface temperature on interannual and longer timescales (1979–2006). Journal of Geophysical Research: Atmospheres 113, (2008).

42. Halali, S. et al. Predictability of temporal variation in climate and the evolution of seasonal polyphenism in tropical butterfly communities. Journal of Evolutionary Biology 34, 1362–1375 (2021).

43. Subash, N. & Sikka, A. K. Trend analysis of rainfall and temperature and its relationship over India. Theor Appl Climatol 117, 449–462 (2014).

44. De Jong, M., Kesbeke, F., Brakefield, P. & Zwaan, B. Geographic variation in thermal plasticity of life history and wing pattern in Bicyclus anynana. Climate Research - CLIMATE RES 43, 91–102 (2010).

45. Nokelainen, O., van Bergen, E., Ripley, B. S. & Brakefield, P. M. Adaptation of a tropical butterfly to a temperate climate. Biological Journal of the Linnean Society 123, 279–289 (2018).

46. Hidalgo, S. & Chiu, J. C. Integration of photoperiodic and temperature cues by the circadian clock to regulate insect seasonal adaptations. Journal of Comparative Physiology. A, Neuroethology, Sensory, Neural, and Behavioral Physiology 210, 585 (2023).

47. Molleman, F. et al. Larval growth rate is not a major determinant of adult wing shape and eyespot size in the seasonally polyphenic butterfly Melanitis leda. PeerJ 12, e18295 (2024).

48. Singh, P. et al. Complex multi-trait responses to multivariate environmental cues in a seasonal butterfly. Evol Ecol 34, 713–734 (2020).

49. van Bergen, E., Atencio, G., Saastamoinen, M. & Beldade, P. Thermal plasticity in protective wing pigmentation is modulated by genotype and food availability in an insect model of seasonal polyphenism. Functional Ecology 38, 1765–1778 (2024).

50. Latorre, B. The Geographic Mosaic of Wolbachia Infection in Melanitis leda Butterfly Populations. Dissertations and Theses (2018).

51. Bujan, J., Yanoviak, S. P. & Kaspari, M. Desiccation resistance in tropical insects: causes and mechanisms underlying variability in a Panama ant community. Ecol Evol 6, 6282–6291 (2016).

52. Chown, S. L. & Gaston, K. J. Body size variation in insects: a macroecological perspective. Biological Reviews 85, 139–169 (2010).

53. Kutrup, B., Bülbül, U. & Özdemir, N. Effects of the ecological conditions on morphological variations of the Green toad, Bufo viridis, in Turkey. Ecological Research 21, 208–214 (2006).

54. Cerdá, X. & Retana, J. Alternative strategies by thermophilic ants to cope with extreme heat: individual versus colony level traits. Oikos 89, 155–163 (2000).

55. Krupp, J. J., Nayal, K., Wong, A., Millar, J. G. & Levine, J. D. Desiccation resistance is an adaptive life-history trait dependent upon cuticular hydrocarbons, and influenced by mating status and temperature in *D. melanogaster*. Journal of Insect Physiology 121, 103990 (2020).

56. Kleynhans, E. & Terblanche, J. S. The evolution of water balance in Glossina (Diptera: Glossinidae): correlations with climate. Biology Letters 5, 93–96 (2008).

57. Fick, S. E. & Hijmans, R. J. WorldClim 2: new 1-km spatial resolution climate surfaces for global land areas. International Journal of Climatology 37, 4302–4315 (2017).

58. NASA Langley Research Center. Prediction Of Worldwide Energy Resources (POWER) Project, Hourly Service. NASA Earth Science Division (2025).

59. Samatha, B. Ecobiology of the Common Evening Brown, Melanitis leda Linn. (Lepidoptera: Rhopalocera: Nymphalidae: Satyrinae). IJARST 617–622 (2016) doi:10.62226/ijarst20160278.

60. Dhamodharan, V., Gandhi, M. S. & Manjula, V. Long-term rainfall trend of Kerala, Tamil Nadu and Pondicherry using departure analysis. 8, 152–157 (2015).

61. Schneider, C. A., Rasband, W. S. & Eliceiri, K. W. NIH Image to ImageJ: 25 years of image analysis. Nat Methods 9, 671–675 (2012).

62. R Core Team. R: A language and environment for statistical computing. (2021).

63. RStudio Team. RStudio:Integrated Development for R. (2022).

64. Rigby, R. A. & Stasinopoulos, D. M. Generalized additive models for location, scale and shape. Journal of the Royal Statistical Society: Series C (Applied Statistics) 54, 507–554 (2005).

65. Gordon, A. D., Johnson, S. E. & Louis Jr., E. E. Females are the ecological sex: Sex-specific body mass ecogeography in wild sifaka populations (Propithecus spp.). American Journal of Physical Anthropology 151, 77–87 (2013).

66. Moura-Campos, D., Chung, M.-H. J., Lawrence, E., Jennions, M. D. & Head, M. L. Temperature-dependent differences in male and female life history responses to a period of food limitation during development. Journal of Animal Ecology 94, 1076–1087 (2025).

67. Stasinopoulos, M., Rigby, B. & Akantziliotou, C. Instructions on how to use the gamlss package in R Second Edition. (2008).

68. Taniguchi, M. & Hirukawa, J. Generalized information criterion. Journal of Time Series Analysis 33, 287–297 (2012).

69. Lenth, R., Singmann, H., Love, J., Buerkner, P. & Herve, M. emmeans: Estimated Marginal Means, aka Least-Squares Means. (2020).

70. Sekar, S. & Karanth, P. Flying between Sky Islands: The Effect of Naturally Fragmented Habitat on Butterfly Population Structure. PloS one 8, e71573 (2013).

71. Mallick, S. et al. Seasonal plasticity in sympatric Bicyclus butterflies in a tropical forest where temperature does not predict rainfall. Biotropica (2024) doi:10.1111/btp.13365.

72. van Bergen, E. et al. Conserved patterns of integrated developmental plasticity in a group of polyphenic tropical butterflies. BMC Evolutionary Biology 17, 59 (2017).

73. Kooi, R. E., Brakefield, P. M. & Rossie, W. E. M. T. Effects of food plant on phenotypic plasticity in the tropical butterfly Bicyclus anynana. Entomologia Experimentalis et Applicata (Netherlands) (1996).

74. Chia, S. Y. & Lim, M. W. A critical review on the influence of humidity for plant growth forecasting. IOP Conf. Ser.: Mater. Sci. Eng. 1257, 012001 (2022).

75. Wu, Z., Dijkstra, P., Koch, G. W., Peñuelas, J. & Hungate, B. A. Responses of terrestrial ecosystems to temperature and precipitation change: a meta-analysis of experimental manipulation. Global Change Biology 17, 927–942 (2011).

76. Beldade, P. & Monteiro, A. Eco-evo-devo advances with butterfly eyespots. Curr Opin Genet Dev 69, 6–13 (2021).

77. Faull, K. J. & Williams, C. R. Intraspecific variation in desiccation survival time of Aedes aegypti (L.) mosquito eggs of Australian origin. J Vector Ecol 40, 292–300 (2015).

78. Fox, C. W. & Czesak, M. E. Evolutionary Ecology of Progeny Size in Arthropods. Annual Review of Entomology 45, 341–369 (2000).

79. Maeno, K. O., et al. Desiccated desert locust embryos reserve yolk as a “lunch box” for posthatching survival. PNAS Nexus 4, pgaf132 (2025).

80. Pijpe, J., Brakefield, P. M. & Zwaan, B. J. Increased Life Span in a Polyphenic Butterfly Artificially Selected for Starvation Resistance. The American Naturalist 171, 81–90 (2008).

81. Ramos-Robles, M. I., Romero, K. L. R., Burgos-Solorio, A. & Aguilar-Dorantes, K. M. The effect of tropical dry forest seasonality on the diversity of insects associated with ferns. Rodriguésia 74, e00362023 (2023).

82. Gershman, S. N., Miller, O. G. & Hamilton, I. M. Causes and consequences of variation in development time in a field cricket. J Evol Biol 35, 299–310 (2022).

83. Semsar-kazerouni, M., Siepel, H. & Verberk, W. C. E. P. Influence of photoperiod on thermal responses in body size, growth and development in Lycaena phlaeas (Lepidoptera: Lycaenidae). Current Research in Insect Science 2, 100034 (2022).

