## Supplementary file for "Local adaptations in wing-pattern and life history trait plasticity in a butterfly: humidity as a cue where temperature is unreliable"

**Table S1. AIC-based model selection results for female relative forewing area. The initial model included all fixed effects. Candidate models were generated by performing a model search utilizing stepGAIC's scope option which sequentially adds or removes predictors, including interaction terms, to identify the model with the lowest AIC. The final selected model is shown in bold.**

| Start model<br>Female relative forewing area ~ TREATMENT + POPULATION AIC = -1121.4 | | | |
|---|---|---|---|
| **Model** | Df | AIC | **ΔAIC** |
| **+ TREATMENT: POPULATION** | 4 | **-1128.8** | **0** |
| – POPULATION | 2 | -1110.7 | 18.1 |
| – TREATMENT | 2 | -1100.0 | 28.8 |

**Table S2. AIC-based model selection results for male relative forewing area. The initial model included all fixed effects. Candidate models were generated by performing a model search utilizing stepGAIC's scope option which sequentially adds or removes predictors, including interaction terms, to identify the model with the lowest AIC. The final selected model is shown in bold.**

| Start model<br>Male relative forewing area ~ TREATMENT + POPULATION AIC = -1210.82 | | | |
|---|---|---|---|
| **Model** | Df | AIC | **ΔAIC** |
| **+ TREATMENT: POPULATION** | 4 | **-1215.2** | **0** |
| – POPULATION | 2 | -1206.3 | 8.9 |
| – TREATMENT | 2 | -1195.8 | 19.4 |

**Table S3. AIC-based model selection results for female relative hindwing area. The initial model included all fixed effects. Candidate models were generated by performing a model search utilizing stepGAIC's scope option which sequentially adds or removes predictors, including interaction terms, to identify the model with the lowest AIC. The final selected model is shown in bold.**

| Start model<br>Female relative hindwing area ~ TREATMENT + POPULATION AIC = -1010.43 | | | |
|---|---|---|---|
| **Model** | Df | AIC | **ΔAIC** |
| **+ TREATMENT: POPULATION** | 4 | **-1017.54** | **0** |
| – POPULATION | 2 | -997.2 | 20.3 |
| – TREATMENT | 2 | -992.68 | 24.8 |

**Table S4. AIC-based model selection results for male relative hindwing area. The initial model included all fixed effects. Candidate models were generated by performing a model search utilizing stepGAIC's scope option which sequentially adds or removes predictors, including interaction terms, to identify the model with the lowest AIC. The final selected model is shown in bold.**

| Start model<br>Male relative hindwing area ~ TREATMENT + POPULATION AIC = -1090.87 | | | |
|---|---|---|---|
| **Model** | Df | AIC | **ΔAIC** |
| **+ TREATMENT: POPULATION** | 4 | **-1098** | **0** |
| – POPULATION | 2 | -1088.1 | 9.9 |
| – TREATMENT | 2 | -1084.1 | 13.9 |

**Table S5. AIC-based model selection results for female pupal weight. The initial model included all fixed effects. Candidate models were generated by performing a model search utilizing stepGAIC's scope option which sequentially adds or removes predictors, including interaction terms, to identify the model with the lowest AIC. The final selected model is shown in bold.**

| Start model<br>**Female pupal weight ~ TREATMENT + POPULATION AIC = -765.71** | | | |
|---|---|---|---|
| **Model** | Df | AIC | **ΔAIC** |
| + TREATMENT: POPULATION | 4 | -761.01 | 4.70 |
| – POPULATION | 2 | -753.98 | 11.73 |
| – TREATMENT | 2 | -763.14 | 2.57 |

**Table S6. AIC-based model selection results for male pupal weight. The initial model included all fixed effects. Candidate models were generated by performing a model search utilizing stepGAIC's scope option which sequentially adds or removes predictors, including interaction terms, to identify the model with the lowest AIC. The final selected model is shown in bold.**

| Start model<br>Male pupal weight ~ TREATMENT + POPULATION AIC = -753.51 | | | |
|---|---|---|---|
| **Model** | Df | AIC | **ΔAIC** |
| + TREATMENT: POPULATION | 4 | -747.71 | 6.16 |
| – POPULATION | 2 | -750.32 | 3.55 |
| **– TREATMENT** | **2** | **-753.87** | **0** |

**Table S7. AIC-based model selection results for female developmental time. The initial model included all fixed effects. Candidate models were generated by performing a model search utilizing stepGAIC's scope option which sequentially adds or removes predictors, including interaction terms, to identify the model with the lowest AIC. The final selected model is shown in bold.**

| Start model<br>Female developmental time ~ TREATMENT + POPULATION AIC = -1122.93 | | | |
|---|---|---|---|
| **Model** | Df | AIC | **ΔAIC** |
| **+ TREATMENT: POPULATION** | **4** | **-1139.4** | **0** |
| – POPULATION | 2 | -1115.6 | 23.80 |
| – TREATMENT | 2 | -1096.4 | 43 |

**Table S8. AIC-based model selection results for male relative developmental time. The initial model included all fixed effects. Candidate models were generated by performing a model search utilizing stepGAIC's scope option which sequentially adds or removes predictors, including interaction terms, to identify the model with the lowest AIC. The final selected model is shown in bold.**

| Start model<br>Male developmental time ~ TREATMENT + POPULATION AIC = -1110.15 | | | |
|---|---|---|---|
| **Model** | Df | AIC | **ΔAIC** |
| **+ TREATMENT: POPULATION** | **4** | **-1136.8** | **0** |
| – POPULATION | 2 | -1084.1 | 52.7 |
| – TREATMENT | 2 | -1070.1 | 66.7 |

**Effects of treatment on pupal mass within populations.**

Among females, in post hoc comparisons show that pupal weight did not differ significantly across treatments in Coimbatore and Vithura populations (all $p > 0.05$; S1). In contrast, Tirunelveli females showed a significant difference between treatments, with pupal weight being higher at 25°C - 85%RH than at 27°C - 85%RH ($E = 0.1606$, $Z = 2.677$, $P = 0.0203$). In males, no pairwise treatment comparisons within population were significant.

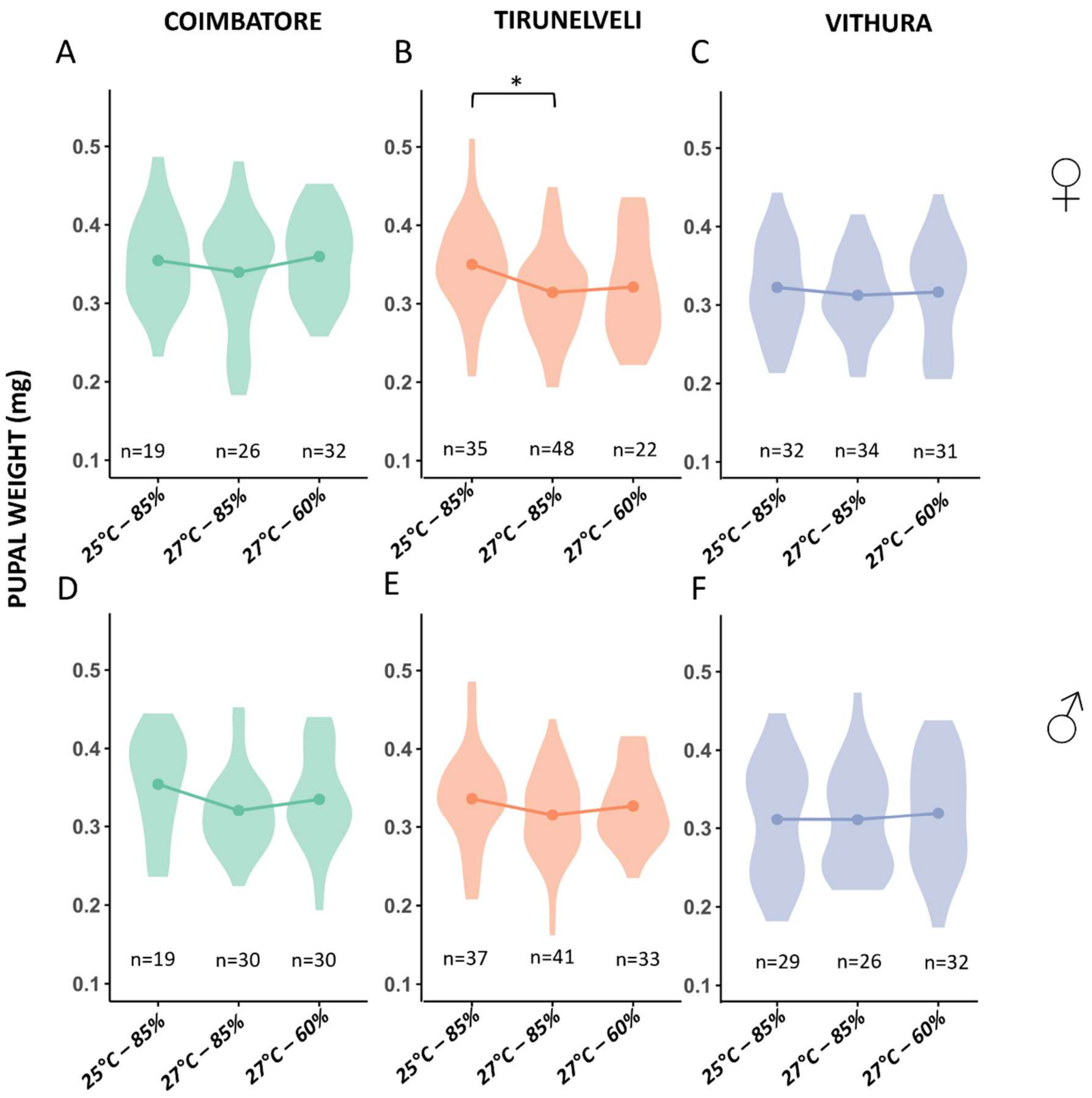

**Figure S1.** Pupal weight across temperature and humidity treatments in *Melanitis leda* populations from Coimbatore, Tirunelveli, and Vithura. Violin plots show the distribution of pupal weight under three environmental conditions: *25 °C – 85%RH*, *27 °C – 85%RH*, and *27 °C – 60%RH*. Data are shown separately for females (A - C) and males (D - F). Points indicate group means. Asterisks denote statistically significant differences between treatment groups ($p < 0.05$ based on *post hoc* pairwise comparisons.

**Effects of treatment on developmental time within populations**

Among females, within populations, no significant treatment-related differences in developmental time were observed in Coimbatore (all $p > 0.05$). In Tirunelveli, developmental time was significantly longer at 25°C - 85%RH than in 27°C - 85%RH ($E = 0.1432$, $Z = 3.517$, $P = 0.0013$), while the other comparisons were not significant. In Vithura, developmental time was significantly longer at 25°C - 85%RH than in 27°C - 85%RH ($E = 0.1482$, $Z = 3.286$, $P = 0.0029$), and significantly longer at 27°C - 60%RH than in 25°C - 85%RH ($E = 0.1493$, $Z = 3.368$, $P = 0.0022$) and 27°C - 85%RH ($E = 0.2975$, $Z = 6.670$, $P < 0.0001$). In males, no significant treatment-related differences in developmental time were observed in Coimbatore (all $p > 0.05$). In Tirunelveli, developmental time was significantly longer at 25°C - 85%RH than in both 27°C - 85%RH ($E = 0.1514$, $Z = 3.642$, $P = 0.0008$) and 27°C - 60%RH ($E = 0.1410$, $Z = 3.209$, $P = 0.0038$), while no difference was detected between the two 27°C treatments. In Vithura males, developmental time was significantly longer at 25°C - 85%RH than in 27°C – 85% ($E = 0.3001$, $Z = 6.079$, $P < 0.0001$), and significantly shorter at 27°C - 85%RH than in 27°C - 60%RH ($E = 0.3807$, $Z = 7.936$, $P < 0.0001$).

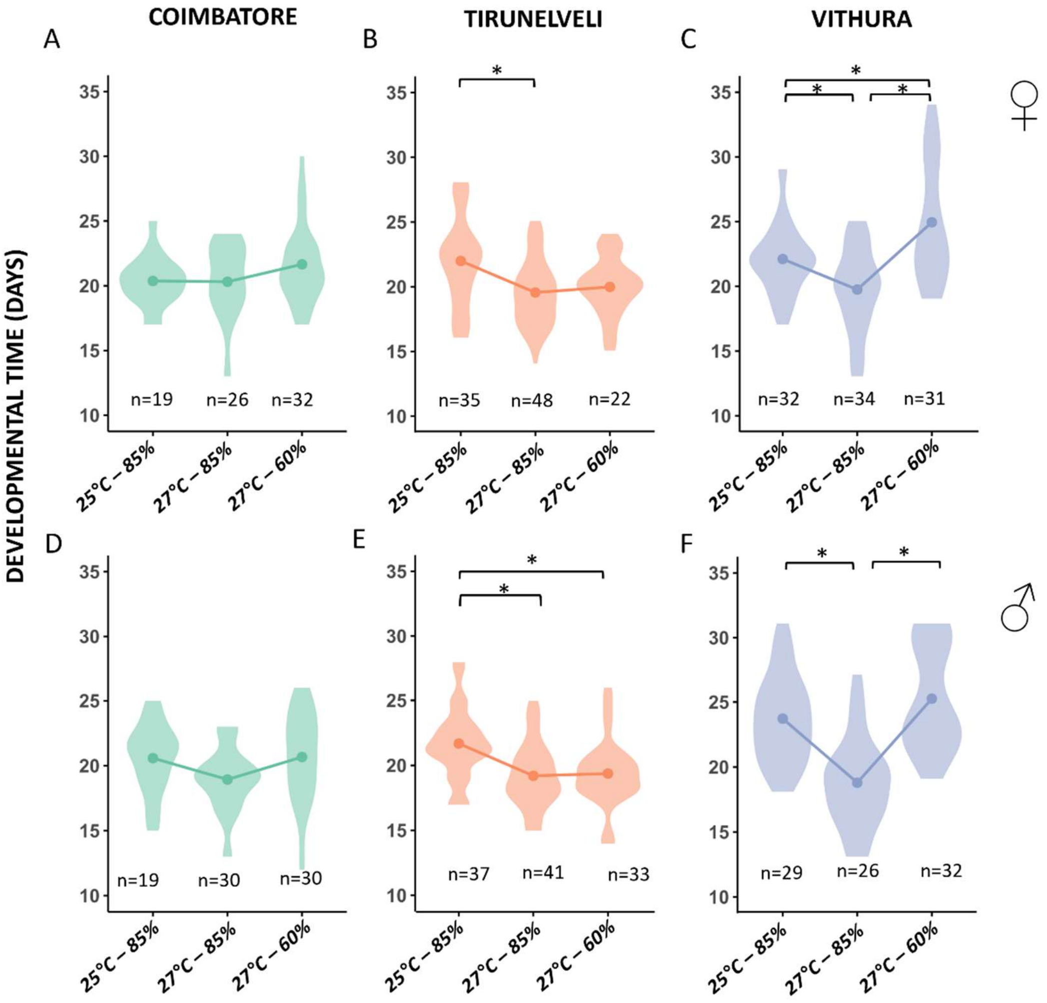

**Figure S2.** Developmental time across temperature and humidity treatments in *Melanitis leda* populations from Coimbatore, Tirunelveli, and Vithura. Violin plots show the distribution of developmental time under three environmental conditions: *25 °C – 85%RH*, *27 °C – 85%RH*, and *27 °C – 60%RH*. Data are shown separately for females (A - C) and males (D - F). Points indicate group means. Asterisks denote statistically significant differences between treatment groups ($p < 0.05$ based on *post hoc* pairwise comparisons).
